# Cancer Stemness Online: A resource for investigating cancer stemness and associations with immune response

**DOI:** 10.1101/2024.03.14.585118

**Authors:** Weiwei Zhou, Minghai Su, Tiantongfei Jiang, Yunjin Xie, Jingyi Shi, Yingying Ma, Kang Xu, Gang Xu, Yongsheng Li, Juan Xu

**Author notes:** These authors contributed equally to this work (Weiwei Zhou, Minghai Su, Tiantongfei Jiang). **Corresponding authors:** Yongsheng Li, Juan Xu.

## Abstract

Cancer progression involves the gradual loss of a differentiated phenotype and acquisition of progenitor and stem-cell-like features, which are potential culprit in immunotherapy resistance. Although the state-of-art predictive computational methods have facilitated predicting the cancer stemness, currently there is no efficient resource that can meet various requirements of usage. Here, we presented the Cancer Stemness Online, an integrated resource for efficiently scoring cancer stemness potential at bulk and single-cell level. The resource integrates 8 robust predictive algorithms as well as 27 signature gene sets associated with cancer stemness for predicting the stemness scores. Downstream analyses were performed from five different aspects, including identifying the signature genes of cancer stemness, exploring the association with cancer hallmarks, cellular states, immune response and communication with immune cells, investigating the contributions for patient survival and the robustness analysis of cancer stemness among different methods. Moreover, the pre-calculated cancer stemness atlas for more than 40 cancer types can be accessed by users. Both the tables and diverse visualization for the analytical results are available for download. Together, Cancer Stemness Online is a powerful resource for scoring cancer stemness and going deeper and wider in the downstream functional interpretation, including immune response as well as cancer hallmark. Cancer Stemness Online is freely accessible at http://bio-bigdata.hrbmu.edu.cn/CancerStemnessOnline.

## Introduction

Although numerous therapeutic modalities have been developed to treat cancer, such as surgery, radiation, chemotherapy and immunotherapy, the risk of cancer recurrence remains high [1]. Cancer progression involves the gradual loss of a differentiated phenotype and acquisition of progenitor and stem-cell-like features [2, 3]. The existence of cancer stem cells (CSCs) has been reported in various cancer types [4]. Cancer stemness has also been reported to be the potential culprit in immunotherapy resistance [5, 6]. A convenient platform providing the markers of cancer stemness and stemness index of patients or cancer cells is critical to understand the potential molecular mechanism and develop useful therapy.

Recently, the state-of-art predictive computational methods facilitate to assess the degree of cancer stemness. The majority of methods mainly based on bulk or single-cell transcriptomes to evaluate the stemness of patients or cancer cells. Briefly, these methods can be classified into unsupervised and supervised methods. For example, the commonly used method was single-sample gene set enrichment analysis (ssGSEA) [7], which estimated the stemness score based on the expressions of collected stemness-related gene signatures. Moreover, CytoTRACE was recently developed to predict the differentiation and developmental potential of single cell by assessing the number of detectably expressed genes per cell [8]. Other tools, such as SLICE [9] and SCENT [10] allow researchers to quantify stemness by entropy analysis. StemID [11] assesses stemness of cell types within a population by utilizing tree topology and transcriptome composition.

On the other hand, numerous supervised methods were also developed to estimate the stemness. mRNAsi is a widely used transcriptome stemness index to evaluate the stemness based on the one-class logistic regression machine learning algorithm [12, 13]. StemnessIndex provides an absolute index to evaluate stemness by comparing the relative expression orderings of the stem cell samples and the normal adult samples from different tissues [13]. In addition, StemSC is a stemness index for single cell [14], which represents the percentage of gene pairs with the same relative expression orderings as the reference of embryonic stem cell samples. All these unsupervised and supervised methods provided valuable tools for estimating the stemness for patients or single cells. However, they were scattered across different literature and are difficult to use for researchers with no programming experience.

Some webservers or databases have been developed to depict cell stemness or collect stem cell-related data. However, the majority of these resources only focus on stem gene sets, without providing stemness of samples from public data directly. For example, SISTEMA [15] collected a large number of human stem cell transcriptome data to display the expression of stem genes under different cell lines, cell types and pathological conditions. StemMapper [16] collected transcriptome data sets of various stem cells. Currently there is no efficient database that can meet various requirements of users.

Therefore, we developed the Cancer Stemness Online (http://bio-bigdata.hrbmu.edu.cn/CancerStemnessOnline/), which is a resource providing the cancer stemness score (CSscore), functional analysis and visualization. To assess the CSscore for bulk or single-cell RNA-seq (scRNA-seq) data, Cancer Stemness Online integrated 5 unsupervised and three supervised methods, which evaluated the differentiation level based on transcriptional complexity or similarity to the reference profiles of stem cells. Basic statistical analysis and additional five advanced analyses modules were provided. Cancer Stemness Online is an online platform that does not require registration. It allows users to upload their data for analysis. It provides multiple visualizations of the results for better understanding the stemness. All charts and tables are available for download. Together, Cancer Stemness Online is a powerful resource for estimating cancer stemness and going deeper and wider in the downstream functional interpretation, including immune response as well as cancer hallmarks.

## Materials and methods

### Collection of cancer stemness gene sets

For collecting the cancer stemness-related gene sets, we queried the studies published in recent years in PubMed with “cancer stem cell” or “stemness” as keywords. In total, we manually curated 2860 articles and recorded 27 canonical cancer stemness gene sets (Table S1). The number of genes ranged from 5 to 1007 in these gene sets. All gene names have mapped to classical gene symbols.

### Quality control

For scRNA-seq data, we removed cells with less than 200 total count and genes expressed in less than 3 cells. Cells with more than 5% mitochondrial gene counts were filtered. For bulk RNA-seq data, samples with no expressed gene were removed. To address the effects of noise and batching of the data, users can use several available tools, such as Seurat [17] and Harmony [18], before uploading it to Cancer Stemness Online.

### Calculation of cancer stemness scores

Cancer Stemness Online collected 8 computational methods to evaluate the stemness potential based on multiple principles. These methods were further categorized into ‘unsupervised’ and ‘supervised’ according to the reference of cancer stem cells. On the other hand, mRNAsi [12], StemnessIndex [13] and GSVA [19] were applied to bulk RNA-seq data. CytoTRACE [20], SLICE [21], SCENT [10], StemSC [14] and GSVA [19] were used for scRNA-seq data. To improve the comparability of results, we carried out 0-1 normalization to all CSscores. In addition to the above methods, the CSscores of scRNA-seq data uploaded by users can also be calculated based on StemID [11].

### Single-cell trajectory analysis

To analyze the cell pseudotime in scRNA-seq data, we performed ‘Monocle 2’ [22], which uses reversed graph embedding to describe multiple fate decisions in a fully unsupervised manner.

### Identification of stemness-related signature

To assess the relevance between CSscores and gene expressions, we calculated the spearman correlation coefficient (SCC). The genes with false discovery rate (FDR) < 0.05 and SCC > 0.5 (default) were identified as cancer stemness-related gene signatures.

### Functional correlations

To investigate the functions of cell types, we first calculated the single sample gene set enrichment analysis (ssGSEA) score for each cell [19]. The cellular states, immune signatures and cancer hallmarks were considered. In addition, we calculated the spearman correlation coefficient between the CSscores and ssGSEA scores. In bulk data, we calculated the infiltration of immune cells in the sample by ‘cibersort’ function [23]. The spearman correlation coefficients between the CSscores and infiltrations of various immune cells were calculated respectively.

### Survival analysis

The clinical information including overall survival and state of samples were uploaded by users. We applied cox proportional hazards regression model to assess the prognosis of all cancer stemness genes, based on their median expression. The K–M survival curves were generated by the ‘survminer’ function with corresponding log-rank P values. For the genes with positive beta of ‘coxph’, we defined them as risky factors, and the negative ones were protective factors.

### Cell–cell communications

To further explore the interactions between cancer stem cells and other cells, we identified the cell–cell communications by iTALK (https://github.com/Coolgenome/iTALK). The integrated ligand-receptor interactions were collected from CellchatDB [24], celltalkDB [25], ICELLNET [26], iTALK, Nichenet [27], singlecellsignalR [28] and one recent study [29]. The union sets of ligand-receptor pairs were integrated in Cancer Stemness Online.

### Database implementation

The frontend of Cancer Stemness Online was built with HTML5, JavaScript, and CSS, and it included the jQuery (v3.3.1), Datatable (1.10.25), ECharts (v5.5.1) and D3 (v7.6.1) plugins. The backend of Cancer Stemness Online was powered by eclipse (MARS.2) and was queried via the Java Server Pages with Apache Tomcat container (v6.0) as the middleware. All data in Cancer Stemness Online were stored and managed using eclipse (MARS.2) and it employed Java and R programs to perform online analyses. Cancer Stemness Online has been tested on several popular web browsers, including Google Chrome, Firefox, and Apple Safari.

## Results

### Overall architecture of Cancer Stemness Online

The purpose of Cancer Stemness Online is to facilitate the prediction of cancer stemness score (CSscore) of tumor cells or samples. The overall design of Cancer Stemness Online was summarized in **Figure 1**. The platform accepts different types of transcriptomes uploaded by users, such as the bulk RNA-seq and scRNA-seq data (Figure 1A). In addition, the users can also upload the clinical data of the patients. The inputted files can be prepared following the format description.

**Figure 1.**
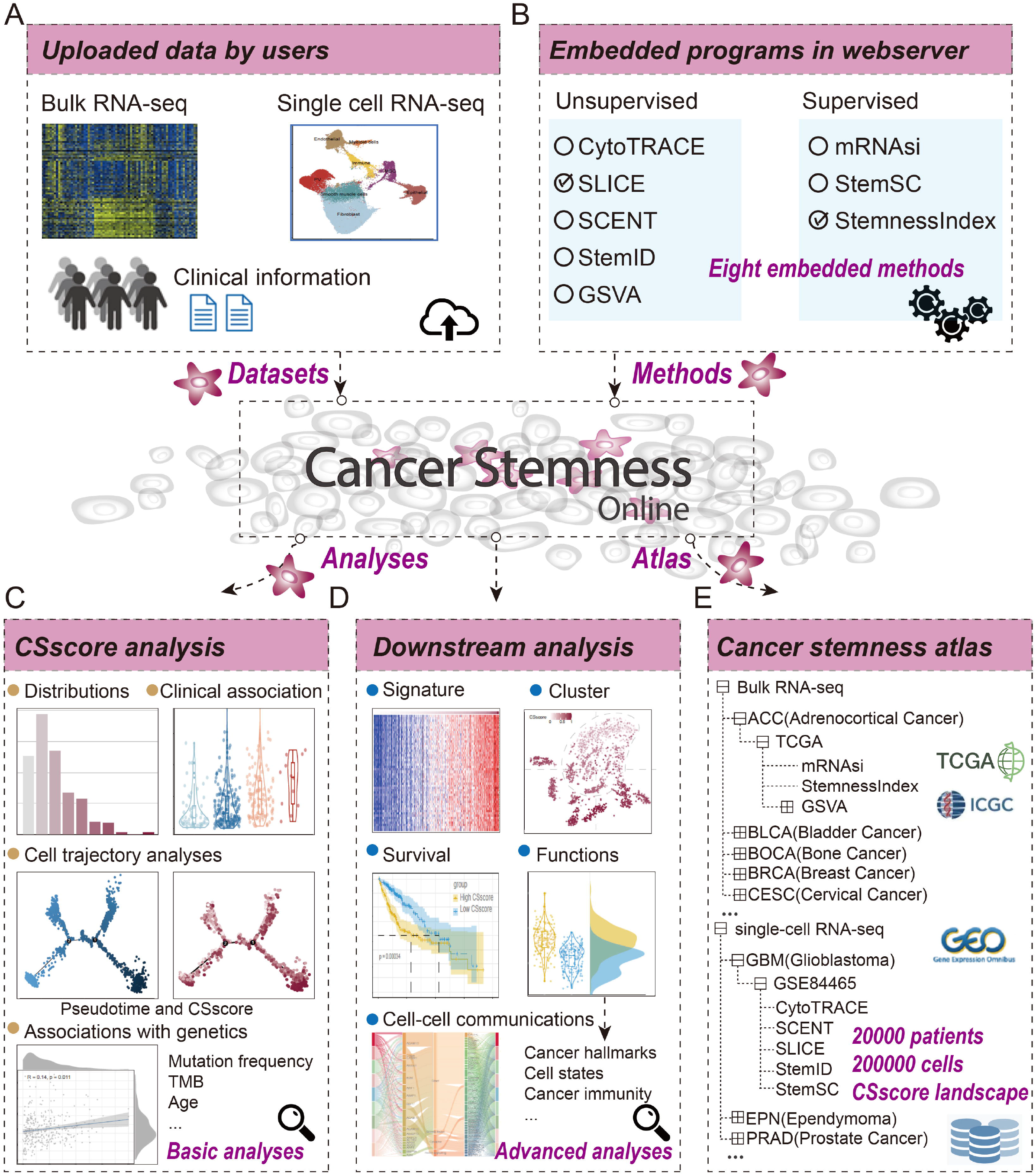
Overall architecture of Cancer Stemness Online. **(A)**, Datasets uploaded by the users, including transcriptomes or clinical information. **(B)**, Robust computational methods embedded in the platform. **(C)**, The basic analysis of cancer stemness in the database. **(D)**, Advanced downstream analysis of the stemness and association with clinical and genetic features. **(E)**, The cancer stemness atlas provided in Cancer Stemness Online.

The platform integrated 8 robust computational algorithms to predict the CSscore for each patient or cell. These methods were classified into five unsupervised and three supervised methods (Figure 1B). To facilitate the selection of the methods, we provided practical guidance from two aspects: By Model Type and By Input Type. Next, the server executes the prediction of CSscore with the selected method. The distribution of CSscores, clinical associations, cell trajectory and associations with genetic features will be returned in the results page (Figure 1C). In addition, the downstream module can identify the gene signatures associated with CSscores, cluster the cells based on expressions of gene signatures, survival analysis, and functional prediction and identify the cell-cell communications (Figure 1D).

Besides the interactive web interface, Cancer Stemness Online also provided flexible ways to access the annotations cancer stemness scores for available cancer transcriptomes projects (Figure 1E), such as TCGA, ICGC and single-cell transcriptomes from published studies. All the analysis results and visualization modules from the resource can be exported as high-quality images and downloaded for further analysis.

### User interface of Cancer Stemness Online

Cancer Stemness Online is an open access online platform for predicting the cancer stemness score for cancer patients or cells. The web interface is freely available and no login is required. The main features of Cancer Stemness Online are the ‘CSscore’ and ‘DownStream’ modules (**Figure 2**). The users can start predicting the CSscore from the ‘GET STARTED’ button in the homepage or from the ‘CSscore’ module. The server allows users to predict the CSscore by selecting from the model type or input type (Figure 2A). In the By Model Type, five unsupervised and three supervised methods can be selected. In the By Input Type, three methods are suitable for bulk transcriptomes and 6 methods for single-cell transcriptomes. The transcriptomes and clinical information of samples can be uploaded and the users can also leave the email information for further retrieving the results from email (Figure 2B).

**Figure 2.**
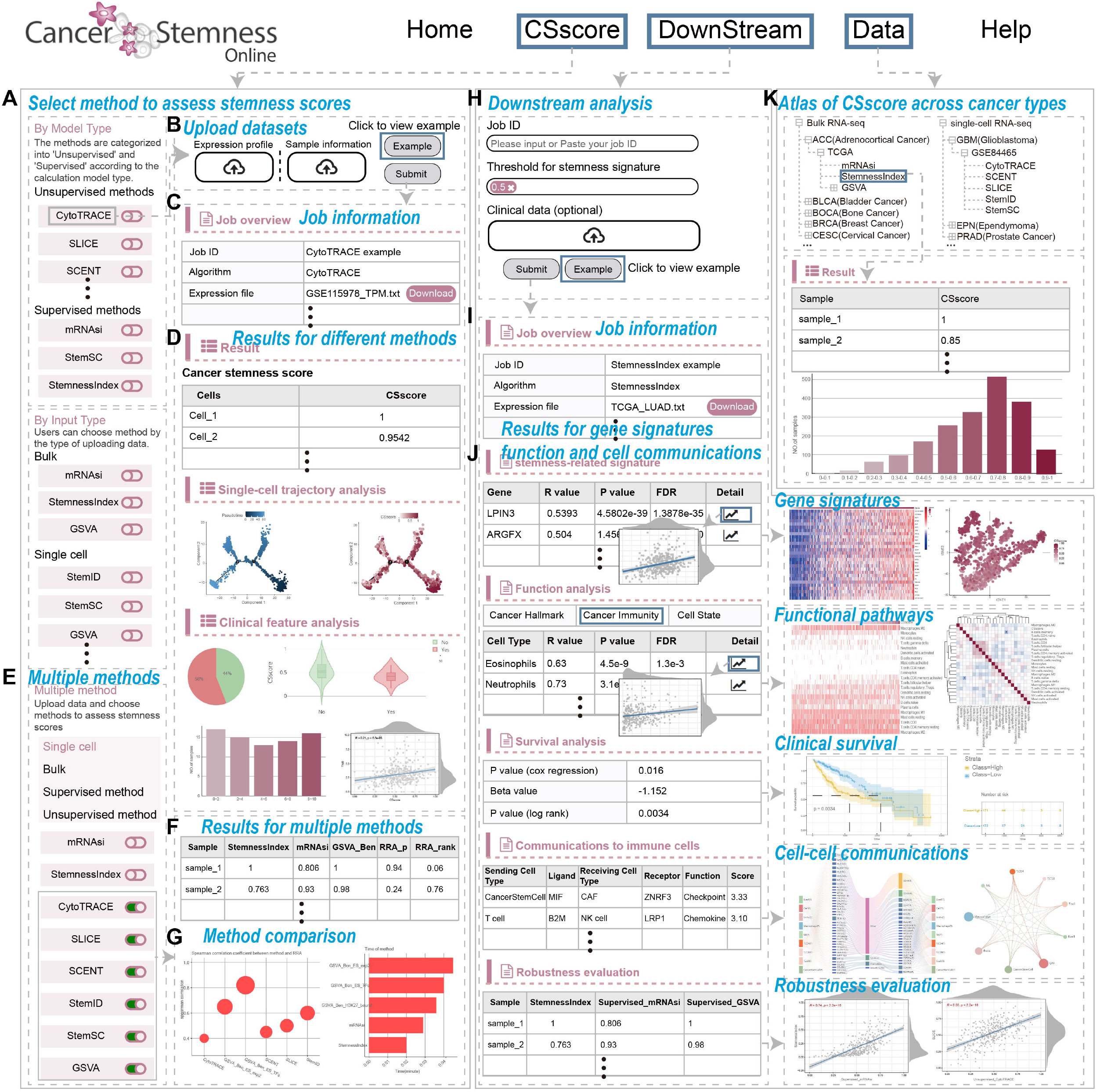
Interactive web interface of Cancer Stemness Online. **(A)**, The methods provided in the platform for users, including unsupervised and supervised methods. **(B)**, The data upload page of the platform. **(C)**, Information of the user submitted job. **(D)**, Results for the basic analysis of CSscores for bulk and single-cell transcriptomes. **(E)**, Screen shot for multiple methods selection page. **(F)**, Results for the multiple methods. **(G)**, Method comparison for different methods. **(H)**, Job submission page for the ‘DownStream’ module. **(I)**, Information for the job submitted by users. **(J)**, Results for the advance analysis, including heat map of gene signatures, functional pathways and immune regulation, clinical survival, cell-cell communications and robustness evaluation. **(K)**, The cancer stemness scores across different cancer types provided in the platform.

The results page first returns the job information, such as the Job ID, algorithm and expression profiles (Figure 2C). The predicted CSscores and associations with clinical features (i.e., grade, tumor mutation burden and treatment) will be provided and visualized in the database (Figure 2D). We also provided a ‘Multiple method’ module in the ‘CSscore’ page, which allows users to select multiple methods and obtained the integrated rank of samples or cells based on the robust rank aggregation (RRA) algorithm (Figure 2E-G). Moreover, the users can perform additional downstream analyses from the ‘DownStream’ module. The users can retrieve the predicted CSscores by inputting the Job ID (Figure 2H). Several parameters can be selected and additional clinical data is optionally uploaded. The new job information will be first provided (Figure 2I) and advanced analysis results will be provided in tables or images (Figure 2J). For example, the genes associated with CSscores will be provided in table and the gene expressions are visualized by heat map. The functional pathways enriched by gene signatures are also provided in table and heat map formats. The clinical survival is performed to evaluate whether the CSscores are associated with survival (Figure 2J). Cell-cell communications and the correlations of CSscores predicted by different methods are also analysed automatically in Cancer Stemness Online. In addition, the predicted CSscores of TCGA, ICGC and single-cell transcriptomes from published studies can be accessed from the ‘Data’ module (Figure 2K). Users can find additional information from the ‘Help’ page.

### Case study 1: Cancer stemness analysis of bulk transcriptomes

To illustrate the various functionalities of Cancer Stemness Online, we first analysed the bulk transcriptomes of hepatocellular carcinoma (HCC) from The Cancer Genome Atlas (TCGA) [30]. We predicted the CSscores for each patient based on the StemnessIndex algorithm (**Figure 3**). We found that the majority of the patients were with low CSscores (Figure 3A), although several patients with high cancer stemness. The server also evaluated the associations between CSscores and clinical features. Cancer patients in high grade were with significantly higher CSscores in HCC (Figure 3B). The CSscores of cancer patients were positively correlated with the number of mutations (Figure 3C, R = 0.14, p = 0.011), which was consistent with previous studies [12, 13, 31]. These results suggested that the CSscore was associated with clinical and genetic features in HCC.

**Figure 3.**
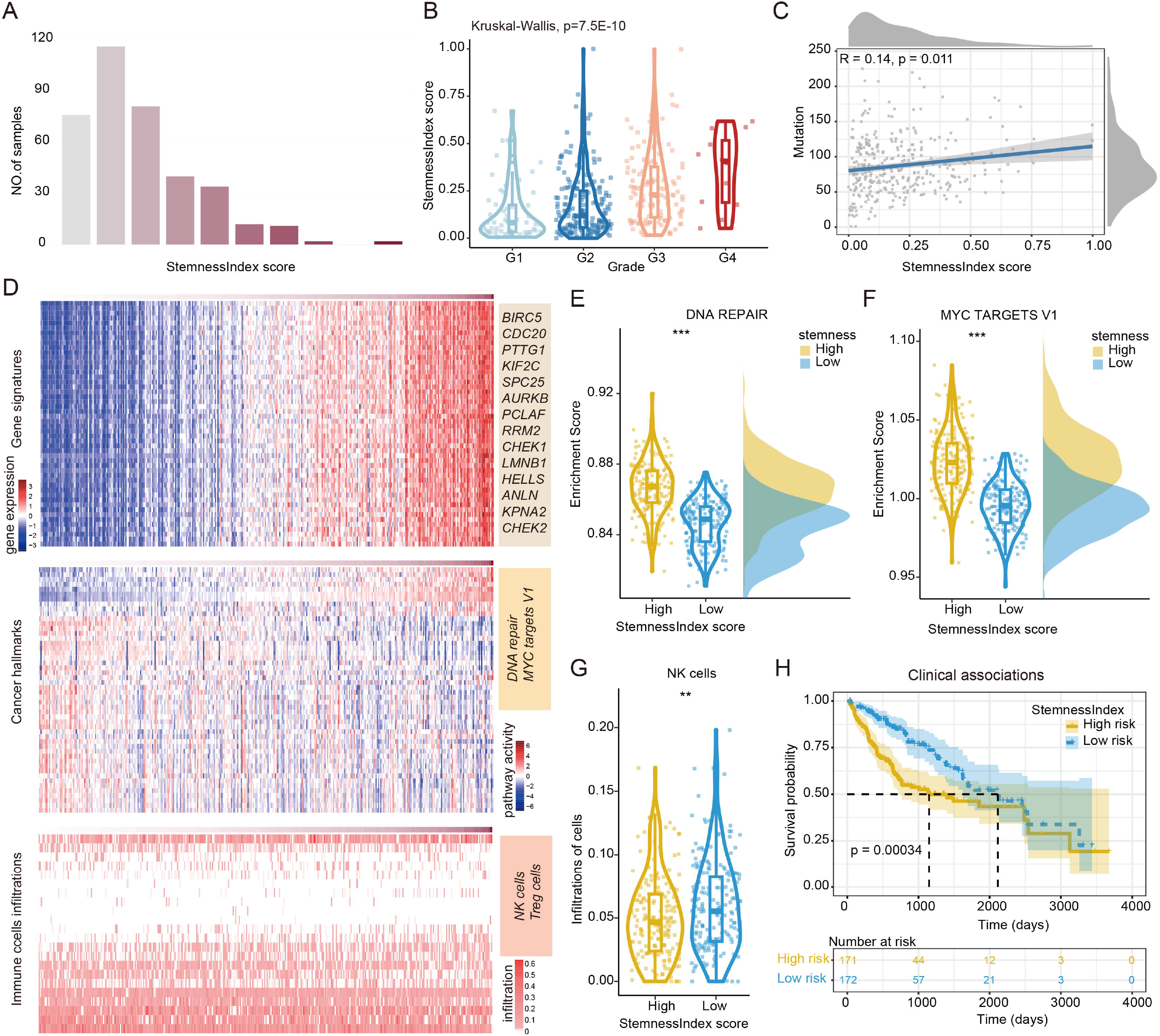
Cancer stemness analysis of bulk transcriptomes in hepatocellular carcinoma. **(A)**, Distributions of CSscores across hepatocellular carcinoma patients. **(B)**, StemnessIndex scores of patients in different grades. **(C)**, Scatter plot showing the correlation between StemnessIndex scores and number of mutations in cancer patients. **(D)**, Heat maps showing the expression of gene signatures, activities of cancer hallmark pathways, and infiltration of immune cells. **(E)**, Box plots showing the enrichment scores of DNA repair pathway in patients with high or low StemnessIndex scores. **(F)**, Box plots showing the enrichment scores of MYC targets V1 in patients with high or low StemnessIndex scores. **(G)**, Box plots showing the infiltration of NK cells in patients with high or low StemnessIndex scores. **(H)**, Kaplan–Meier curve for overall survival of patients with high or low StemnessIndex scores.

Next, we performed advanced analyses based on the ‘DownStream’ module of Cancer Stemness Online. We identified numerous of genes whose expressions were associated with CSscores in HCC (Figure 3D), including BIRC5 [32], CDC20 [33], PTTG1 [34], and KIF2C [35]. Functional analyses revealed that the ‘DNA repair’ and ‘MYC targets V1’ pathways, infiltrations of several immune cells were significantly associated with CSscores of cancer patients (Figure 3D). In particular, cancer patients with high CSscores exhibited significantly higher enrichment scores of ‘DNA repair’ (Figure 3E, p < 0.001) and ‘MYC targets V1’ (Figure 3F, p < 0.001). In addition, there were significantly higher infiltrations of NK cells in cancer patients with low CSscores (Figure 3G, p < 0.01). We next evaluated the survival rates of patients with different CSscores and found that patients with higher stemness exhibited significantly poor survival in HCC (Figure 3H, p = 0.00034, log-rank test). These results suggested that Cancer Stemness Online not only predicted the cancer stemness accurately, but also provided novel insights into the functional pathways and immune regulation in cancer.

### Case study 2: Cancer stemness analysis of single-cell transcriptomes

The development of single-cell sequencing in cancer research has revolutionized our understanding of the biological characteristics within different cancer types [36]. We next analysed the cancer stemness of single-cell transcriptome based on the Cancer Stemness Online server. We obtained the single-cell transcriptome of melanoma from one previous study [37], including 7186 cells from 31 patients. We estimated the CSscores for each cancer cell based on CytoTRACE algorithm embedded in the server (**Figure 4**). We found that large numbers of cells were with higher CSscores in melanoma (Figure 4A). In addition, the pseudotime of cells was estimated by monocle and we found that cells with low pseudotime exhibited significantly higher CSscores (Figure 4B). Immune checkpoint inhibitors (ICI) produce durable responses in some melanoma patients. We found that cells from post treatment were with significantly higher CSscores than those of treatment naive (Figure 4C, p < 2.2E-16), suggesting potential immunotherapy resistance [31].

**Figure 4.**
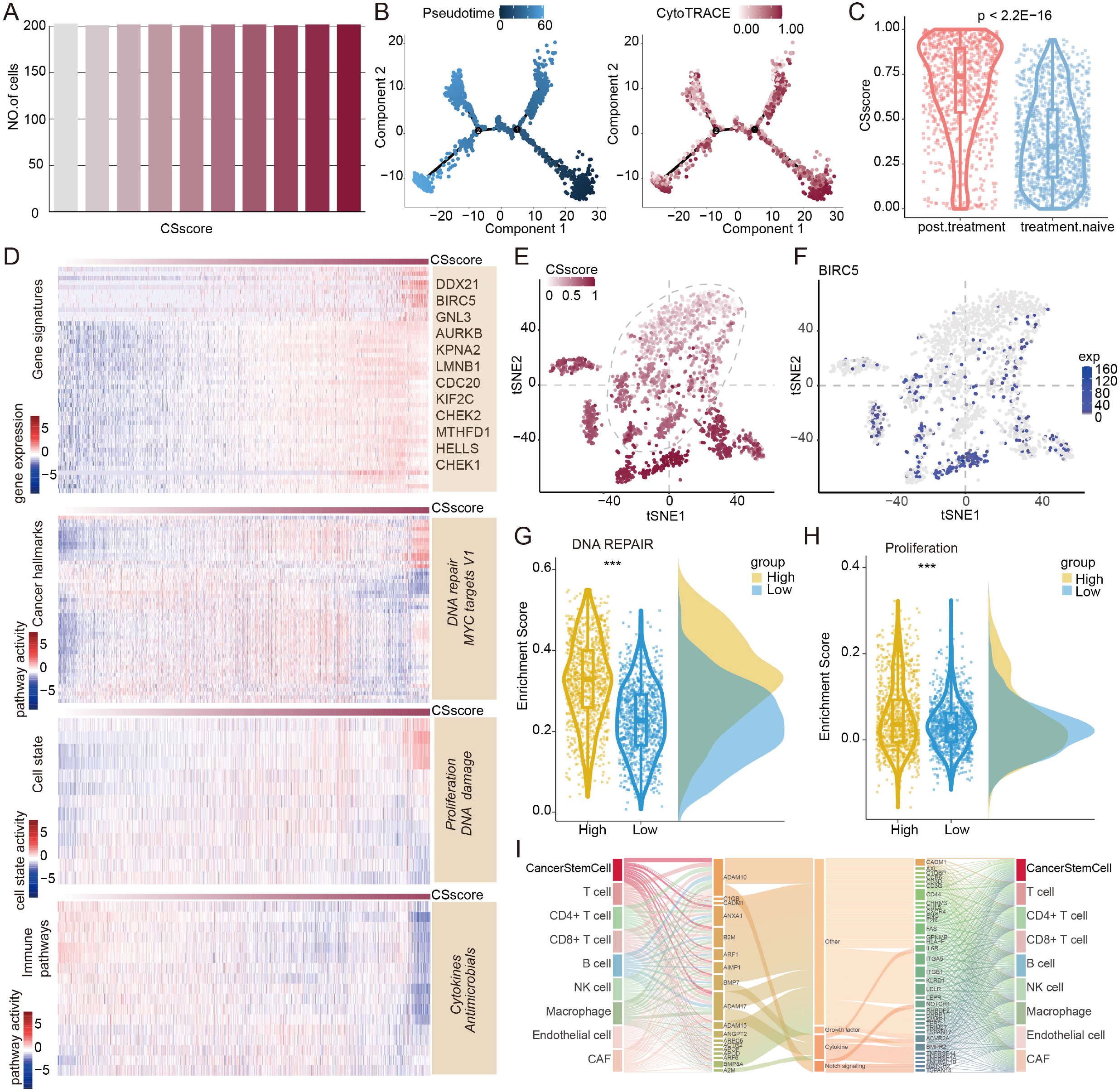
Cancer stemness analysis of single-cell transcriptomes in melanoma. **(A)**, Number of cells with different CSscores. **(B)**, tSNE plot showing the cells with different pseudotime and CytoTRACE scores. **(C)**, Distribution of CSscores for cells from post treatment and naive. **(D)**, Heat maps showing the expressions of gene signatures, activities of cancer hallmark pathways or cell states, and immune pathways. **(E)**, tSNE plot showing the distribution of cells based on expressions of gene signatures. **(F)**, tSNE plot showing the distribution of cells coloured by expression of BIRC5. **(G)**, Box plots showing the enrichment scores of DNA repair pathway in cancer cells with high or low CSscores. **(H)**, Box plots showing the enrichment scores of proliferations in cancer cells with high or low CSscores. **(I)**, Cell-cell communications mediated by ligand-receptor interactions.

In the ‘DownStream’ module, we first identified numerous genes whose expressions were correlated with CSscores (Figure 4D). We found that the expressions of genes can effectively distinguish the cells with higher or lower CSscores (Figure 4E). For example, BIRC5 was highly expressed in cells with higher CSscores (Figure 4F). Functional pathway and immune regulation analyses revealed that cancer cells with high CSscores exhibited significantly enrichment of DNA repair (Figure 4G, p < 0.001) and proliferation (Figure 4H, p < 0.001). These results were consistent with previous observations [31, 38, 39]. We also investigated the cell-cell communications based on the ligand-receptor interactions. We found that cancer stem cells communicated with other immune cells via various ligand-receptor interactions (Figure 4I). In particular, interaction between ADAM10-CD44 helps communication between cancer stem cells and T cells (Figure 4I) [40, 41]. All the analysis results visualized on the web interface were available for download.

## Discussion

Cancer Stemness Online is a useful resource for scoring cancer stemness and associations with immune response, which integrated 8 robust predictive algorithms. The platform supports different types of input transcriptomes and the output of Cancer Stemness Online provided the tables and images for visualization of the CSscores and associations with clinical features. These results benefit the non-computational biologists to explore the cancer stemness. Cancer Stemness Online encompasses not only diverse functionalities, but also user-friendly operations and visually intuitive interfaces. In addition, recent studies have shown that a high stemness profile in cancer is associated with an inferior immunogenic response [42]. Different types of immune cells can be recruited from tumor-associated stem cells [43-45]. Thus, the ‘DownStream’ module in Cancer Stemness Online provided advanced analysis for investigating the functional pathway and immune regulation in the context of cancer stemness. Overall, Cancer Stemness Online is a user-friendly platform to predict the cancer stemness and explore the functional consequence in cancer.

We provided diverse methods to predict the stemness scores for individual sample or cell. To further assist the users selecting the appropriate methods, we first compared the performances of different methods based on both bulk and single cell methods from recent researches [46, 47]. We found that the method ‘StemnessIndex’ might be the most effective for bulk transcriptomes, while ‘CytoTRACE’ might be the most effective one for single-cell transcriptomes (Figure S1). In addition, we have provided a ‘Multiple method’ section module in the ‘CSscore’ page. This module allows users to select multiple methods to predict the stemness scores, and we next obtained the integrated rank of samples or cells based on the robust rank aggregation (RRA) algorithm. The runtimes and correlations between different methods and RRA were provided. Thus, users can integrate the results from multiple methods for downstream analysis.

Nevertheless, there are still rooms to improve in the future. Here are a few areas that we plan to expand in the future version of Cancer Stemness Online. (1) improve the coverage of computational methods and cancer stemness gene sets. Currently, 8 computational methods were integrated in Cancer Stemness Online. We plan to cover newly developed algorithms and cancer stemness gene sets in the near future. (2) expand to cover additional genomes and transcriptomes. The server can only predict the CSscores for human transcriptomes, which should consider in the future working for a wider species. With the development of high throughput sequencing technology, additional cancer transcriptomes will be added in the cancer stemness atlas. (3) include additional annotations. We plan to add more functional annotations, such as more immune cells, more pathways and more immunotherapy information.

Overall, Cancer Stemness Online is a powerful resource for reducing the barrier to analyse the huge transcriptome data that biomedical researchers face and facilitating the identification of association with cancer immunotherapy response for further mechanistic and functional insights.

## Data availability

The web server of Cancer Stemness Online is freely accessible at http://bio-bigdata.hrbmu.edu.cn/CancerStemnessOnline

## Code availability

Code used to perform analyses in this manuscript is available at https://github.com/ComputationalEpigeneticsLab/CancerStemnessOnline.

## CRediT author statement

**Weiwei Zhou:** Conceptualization, Software, Formal analysis, Visualization, Methodology, Writing–original draft. **Minghai Su:** Conceptualization, Software, Formal analysis, Visualization, Methodology, Writing–original draft. **Tiantongfei Jiang:** Conceptualization, Software, Formal analysis, Visualization, Methodology, Writing–original draft. **Yunjin Xie:** Conceptualization, Methodology, Writing–review & editing. **Jingyi Shi:** Software, Methodology, Writing–review & editing. **Yingying Ma:** Conceptualization, Software, Methodology, Writing–review & editing. **Kang Xu:** Conceptualization, Software, Methodology, Writing–review & editing. **Gang Xu:** Conceptualization, Software, Writing– review & editing. **Yongsheng Li:** Conceptualization, Formal analysis, Supervision, Methodology, Project administration, Writing– review & editing. **Juan Xu:** Conceptualization, Formal analysis, Supervision, Methodology, Project administration, Writing– review & editing.

## Acknowledgments

This work was supported by the National Natural Science Foundation of China (32322020, 32170676, 32060152); Natural Science Foundation of Heilongjiang Province (Key Program) (ZD2023C007) and Heilongjiang Touyan Innovation Team Program.

## Supplementary material

**Figure S1. Accuracy of cancer stemness methods**. (A), The correlation between CS scores and differentiation days as calculated by the three bulk methods. (B), The correlation between CS scores and differentiation days as calculated by the six single cell methods. The score was calculated using Spearman’s correlation coefficient.

**Table S1. Stemness marker gene sets used in this study**.

