## Supplementary material for "Cancer Stemness Online: A resource for investigating cancer stemness and associations with immune response": Figure S1

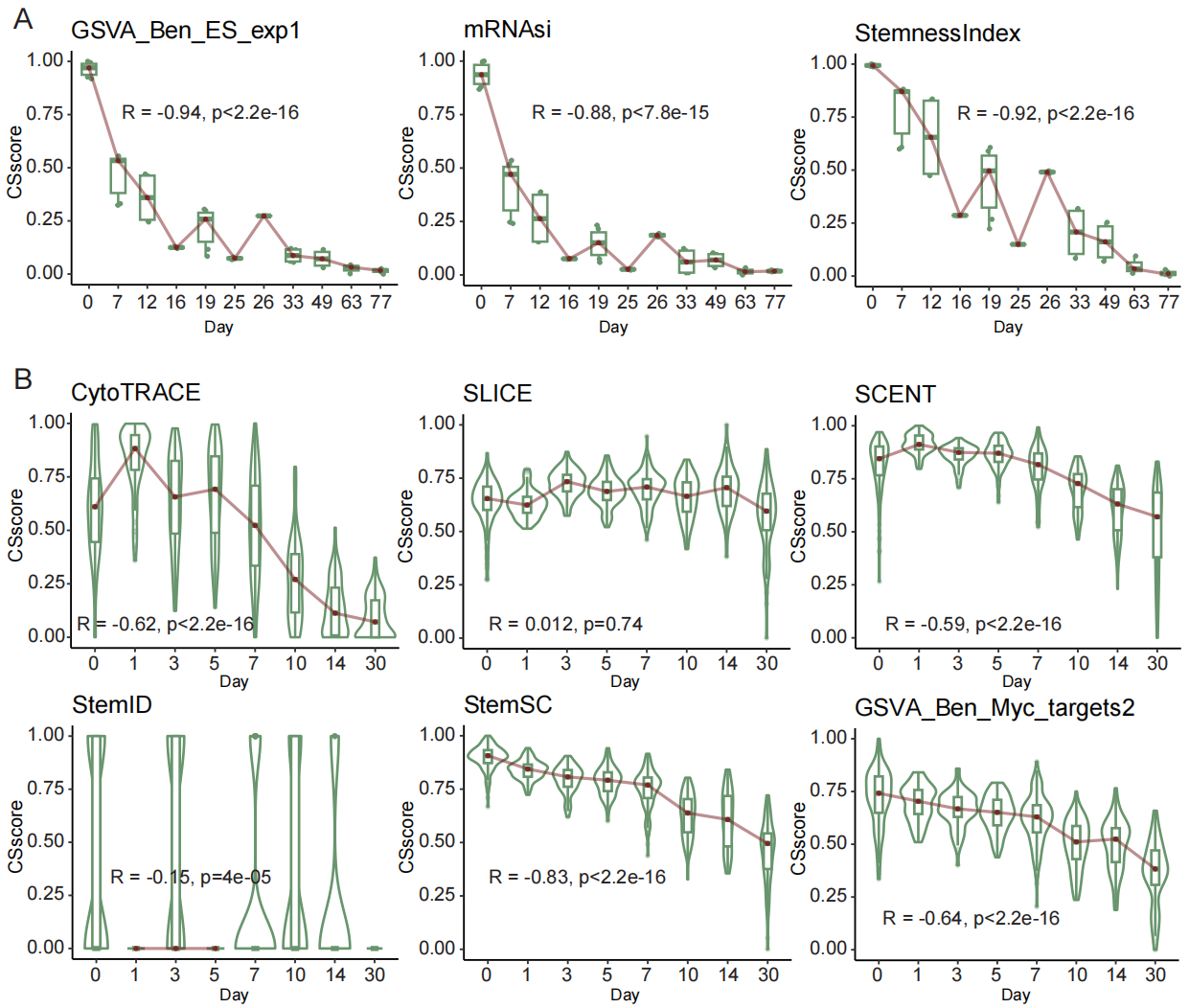

**Fig. S1. Accuracy of cancer stemness methods.** (A), The correlation between CS scores and differentiation days as calculated by the three bulk methods. (B), The correlation between CS scores and differentiation days as calculated by the six single cell methods. The score was calculated using Spearman's correlation coefficient.
